# Characterisation and prion transmission study in mice with genetic reduction of sporadic Creutzfeldt-Jakob Disease risk gene *Stx6*

**DOI:** 10.1101/2023.01.10.523281

**Authors:** Emma Jones, Elizabeth Hill, Jacqueline Linehan, Tamsin Nazari, Adam Caulder, Gemma F Codner, Marie Hutchison, Matthew Mackenzie, Michael Farmer, Thomas Coysh, Michael Wiggins De Oliveira, Huda Al-Doujaily, Malin Sandberg, Emmanuelle Viré, Thomas J Cunningham, Emmanuel A Asante, Sebastian Brandner, John Collinge, Simon Mead

**Author notes:** These authors contributed equally to this work (Emma Jones, Elizabeth Hill). AsanSka University College of Design and Technology, P.O. Box VV 179, Oyibi, Ghana.

## Abstract

Sporadic Creutzfeldt-Jakob disease (sCJD), the most common human prion disease, is thought to occur when the cellular prion protein (PrP^C^) spontaneously misfolds and assembles into prion fibrils, culminating in fatal neurodegeneration. In a genome-wide association study of sCJD, we recently identified risk variants in and around the gene *STX6*, with evidence to suggest a causal increase of *STX6* expression in disease-relevant brain regions. *STX6* encodes syntaxin-6, a SNARE protein primarily involved in early endosome to *trans*-Golgi network retrograde transport. Here we developed and characterised a mouse model with genetic depletion of *Stx6* and investigated a causal role of *Stx6* expression in mouse prion disease through a classical prion transmission study, assessing the impact of homozygous and heterozygous syntaxin-6 knockout on disease incubation periods and prion-related neuropathology. Following inoculation with RML prions, incubation periods in *Stx6*^-/-^ and *Stx6*^+/-^ mice differed by 12 days relative to wildtype. Similarly, in *Stx6*^-/-^ mice, disease incubation periods following inoculation with ME7 prions also differed by 12 days. Histopathological analysis revealed a modest increase in astrogliosis in ME7-inoculated *Stx6*^-/-^ animals and a variable effect of *Stx6* expression on microglia activation, however no differences in neuronal loss, spongiform change or PrP deposition were observed at endpoint. Importantly, *Stx6*^-/-^ mice are viable and fertile with no gross impairments on a range of neurological, biochemical, histological and skeletal structure tests. Our results provide some support for a pathological role of *Stx6* expression in prion disease, which warrants further investigation in the context of prion disease but also other neurodegenerative diseases considering syntaxin-6 appears to have pleiotropic risk effects in progressive supranuclear palsy and Alzheimer’s disease.

**Author Summary:** Sporadic Creutzfeldt-Jakob disease (sCJD), the most common human prion disease, is an invariably fatal disease with no established disease-modifying treatments. The identification of *STX6* as a proposed risk gene for sCJD motivated the generation of a new mouse knockout model, in which we found no grossly deleterious phenotypes. A transmission study in *Stx6^-/-^, Stx6^+/-^* and *Stx6*^+/+^ mice challenged with two prion strains showed reduced syntaxin-6 expression is associated with a modest prolongation of prion disease incubation periods, supporting a pathological role of *Stx6* expression in prion disease pathogenesis. Syntaxin-6 appears to have pleiotropic risk effects across multiple neurodegenerative diseases including progressive supranuclear palsy and Alzheimer’s disease. Thus, this work supports further exploration of the *STX6* susceptibility mechanism, which likely has relevance across multiple neurodegenerative diseases.

## Introduction

Prion diseases are transmissible neurodegenerative conditions of humans and animals with no established treatments. The infectious agent of prion diseases, or prion, comprises amyloid assemblies of misfolded host prion protein in a parallel in-register beta sheet conformation(1, 2). Sporadic Creutzfeldt-Jakob disease (sCJD), the most common human prion disease, invariably results in fatal neurodegeneration. sCJD is thought to result from the spontaneous formation of prions in the body by misfolding and aggregation of the host-encoded prion protein (PrP^C^), triggering a self-perpetuating cycle whereby prion fibrils elongate by recruitment of cellular PrP with subsequent fission producing more infectious particles. Variation in the *PRNP* gene encoding PrP^C^ is a key genetic determinant of sCJD risk and phenotype, however there is now robust evidence for two non-*PRNP* genetic risk factors identified through a genome-wide association study. This study found a genetic region including *STX6* as a novel risk locus in human sCJD(3). Interestingly the *STX6* locus has also been implicated in other neurodegenerative diseases, with the same genetic variants being associated with progressive supranuclear palsy (PSP)(4, 5) and increased syntaxin-6 protein expression being causally associated with Alzheimer’s disease (AD)(6, 7), indicating this protein may play a pleiotropic role in neurodegeneration.

*STX6* encodes syntaxin-6, a member of the SNARE (soluble N-ethylmaleimide-sensitive factor attachment protein receptor) protein family, which is involved in the final step of membrane fusion during vesicle transport. Syntaxin-6 has been shown to be primarily involved in vesicle fusion during retrograde transport between early endosomes and the *trans*-Golgi network(8, 9). Analysis of expression quantitative trait loci (eQTLs) at this site identified *cis-*acting variants which increase *STX6* expression in disease-relevant brain regions, particularly the putamen and caudate nuclei, which are commonly found to be abnormal in diagnostic MRI brain studies in sCJD(10). These and other gene prioritisation analyses identify *STX6* as the causal gene driving the increased sCJD risk at this locus.

Experimental inoculation of mice with prions faithfully recapitulates the neuropathological hallmarks seen in human prion diseases, notably the propagation and deposition of PrP aggregates, spongiform degeneration, neuronal loss and widespread glial cell and immune activation(11, 12). Furthermore, it has been long recognised that prions exist as conformationally distinct strains(13), which alter the disease progression and pathology in mice in a similar manner to human diseases. Prion strains RML and ME7 have been developed, whereby prions originally derived from sheep and goat scrapie have been serially passaged in mice resulting in mouse-adapted prions with high attack rates and well-defined incubation periods in a laboratory setting(11, 14). Mouse bioassay has been routinely used to measure prion infectivity(15, 16), as well as to further understand disease modifiers in a mammalian biological system(17, 18). In a standard experiment, the time from inoculation to prion disease diagnosis (termed “incubation period”) is the defining measure.

To explore a role for *Stx6* expression in prion disease pathogenesis, we conducted a classical prion transmission study in a newly developed mouse model which has a genetic reduction in *Stx6* expression. We show that knockout of syntaxin-6 does not result in a deleterious phenotype but had a 12 day longer incubation period following inoculation with RML or ME7 mouse-adapted scrapie prions, representing an 8% and 7% increase in the incubation periods relative to wildtype controls respectively. *Stx6^+/-^* mice showed an 8% prolongation of incubation periods following RML inoculation, although no extension in the incubation period was seen with ME7-inoculated *Stx6^+/-^* animals. No differences in neuronal loss, spongiform change or PrP deposition were seen between the genotypes, however a variable effect of *Stx6* expression on gliosis was observed at endpoint. This work provides complimentary evidence that altered expression of *Stx6* modifies prion disease pathogenesis and supports the further exploration of the functional role of syntaxin-6 in prion disease and related tauopathies.

## Results

### Generation of *Stx6* knockout C57BL/6N mice

We generated mice with homozygous deletion of *Stx6* (*Stx6*^-/-^) on the C57BL/6N background as part of the International Mouse Phenotyping Consortium(19) (IMPC) via CRISPR/Cas9-mediated deletion of 1808 nucleotides spanning two critical exons (exons 6 and 7 of transcripts *Stx6-201* and *Stx6-209* as depicted in **Fig 1A**, present within all protein coding isoforms) replaced by an 8 nucleotide insertion, generating a premature stop codon. Quantitative immunoblotting of whole brain homogenates demonstrated the loss of the primary protein product at ∼32 kDa in mice with homozygous deletion of *Stx6* (*Stx6*^-/-^), whilst heterozygous mice (*Stx6*^+/-^) expressed ∼50% of the protein compared to wildtype animals (*Stx6*^+/+^) (*Stx6*^-/-^: 0.0757% ± 0.0217 (mean ± SEM); *Stx6*^+/-^: 55.9% ± 0.0436) (**Fig 1B-C**). This was also seen with an N-terminal syntaxin-6 antibody (**Error! Reference source not found.**). Reduced expression of PrP^C^ has previously been shown to extend the incubation period in mice(20). Therefore to determine any differences associated with *Stx6* expression, brain PrP^C^ levels were assessed by enzyme-linked immunosorbent assay (ELISA). This analysis did not reveal any differences in PrP^C^ expression with *Stx6* genotype (*Stx6*^+/+^: 6115 ± 838.3 RFU (mean ± SEM); *Stx6*^+/-^: 5784 ± 1100 RFU; *Stx6*^-/-^: 5834 ± 153 RFU) (**Fig 1D**).

**Fig 1:**
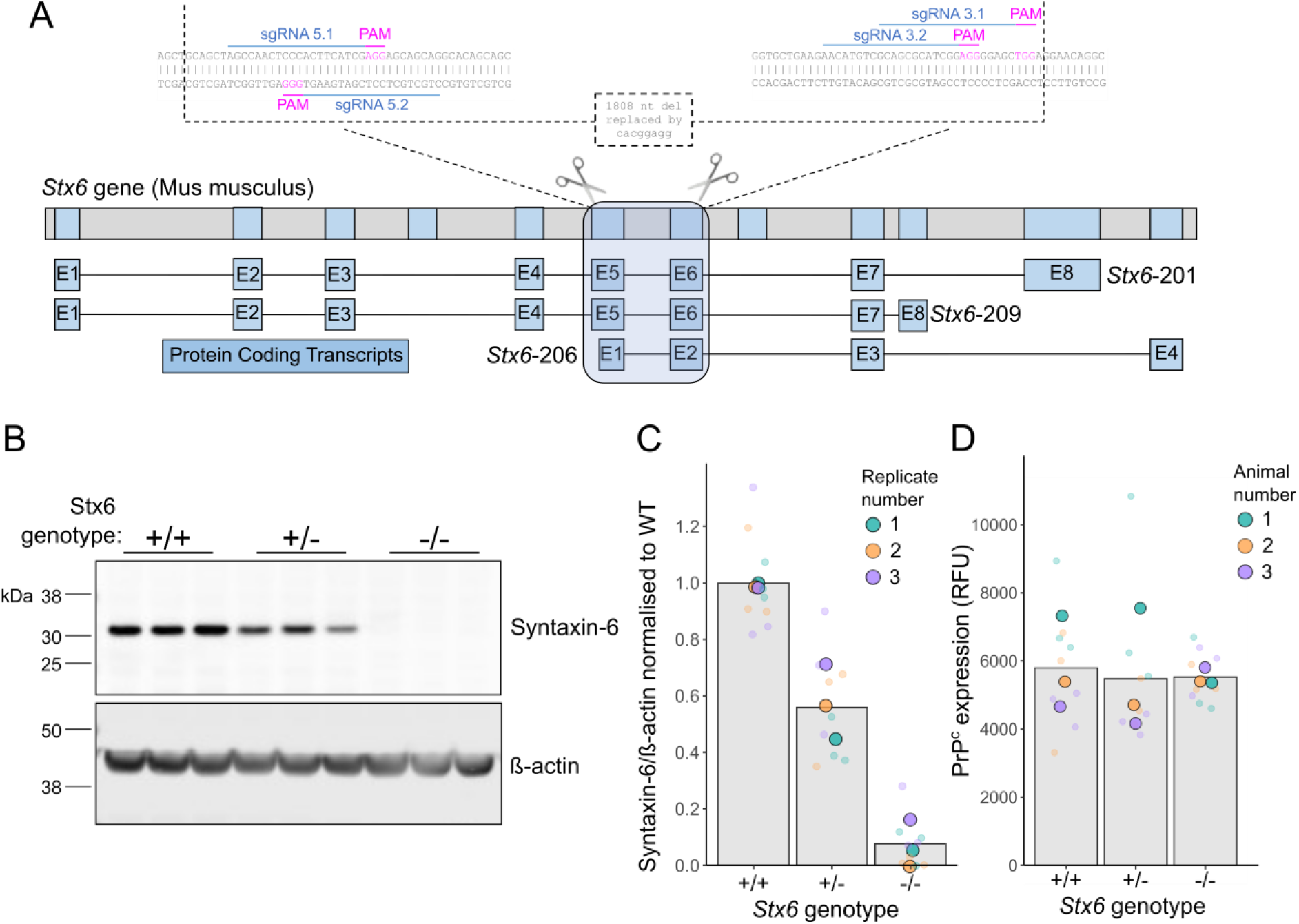
Development and validation of Stx6 knockout mice. **(A)** Strategy for CRISPR/Cas9-mediated knockout of Stx6 in C57BL/6N mice. Four single guide RNAs (sgRNAs) targeting the DNA sequences in exons 5 and 6 of the two main Stx6 protein-coding transcripts (Stx6-201 and Stx6-209), also encompassing the N-terminal portion of the shorter Stx6-206 transcript, were selected for gene editing in zygotes. This resulted in an 1808 nucleotide (nt) deletion replaced by an 8 nt insertion. Top: Full Stx6 gene in the genome. Bottom: protein coding transcripts. Inset: CRISPR/Cas9 genome editing at Stx6 locus. **(B)** Representative quantitative immunoblots with anti-syntaxin 6 and anti-β actin antibodies of whole brain homogenates from Stx6^+/-^, Stx6^-/-^ and Stx6^+/+^ mice and **(C)** quantification of syntaxin-6 intensity relative to β-actin normalised to Stx6^+/+^ control demonstrates loss of primary ∼32 kDa protein isoform in Stx6^-/-^ mice (0.0757% ± 0.0217 (mean ± SEM)) with ∼50% expression (55.9% ± 0.0436) in Stx6^+/-^ mice (n = 3 per group, n = 3 technical replicates). **(D)** Analysis of PrP^C^ expression in whole brain homogenates from Stx6^-/-^, Stx6^+/-^ and Stx6^+/+^ early adult 9 week old mice by ELISA does not indicate any significant differences between lines (Stx6^+/+^: 6115 ± 838.3 relative fluorescent units (RFU); Stx6^+/-^: 5784 ± 1100 RFU; Stx6^-/-^: 5834 ± 153 RFU) (one-way ANOVA on square root transformed data).

### *Stx6*^-/-^ mice are physiologically normal

To determine any gross changes in organ morphology associated with altered *Stx6* expression, immunohistochemical analysis was performed on liver, pancreas, kidney, skeletal muscle and adipose tissue dissected from 4-5 adult mice (4 × *Stx6*^-/-^, 5 × *Stx6*^+/-^, 5 × *Stx6*^+/+^, see Materials and Methods). As shown in **Fig 2**, there were no differences in the appearance and architecture between any tissue from *Stx6*^-/-^, *Stx6*^+/-^ and *Stx6*^+/+^ animals aside from moderate liver steatosis in one *Stx6* null animal, however as this was only found in one animal it is unlikely to be related to the loss of *Stx6* expression (**Error! Reference source not found.**). Furthermore, analysis of skeletal structure by computed tomography (CT) scanning did not reveal any obvious differences in bone or skeleton architecture between genotypes (**Error! Reference source not found.**).

**Fig 2:**
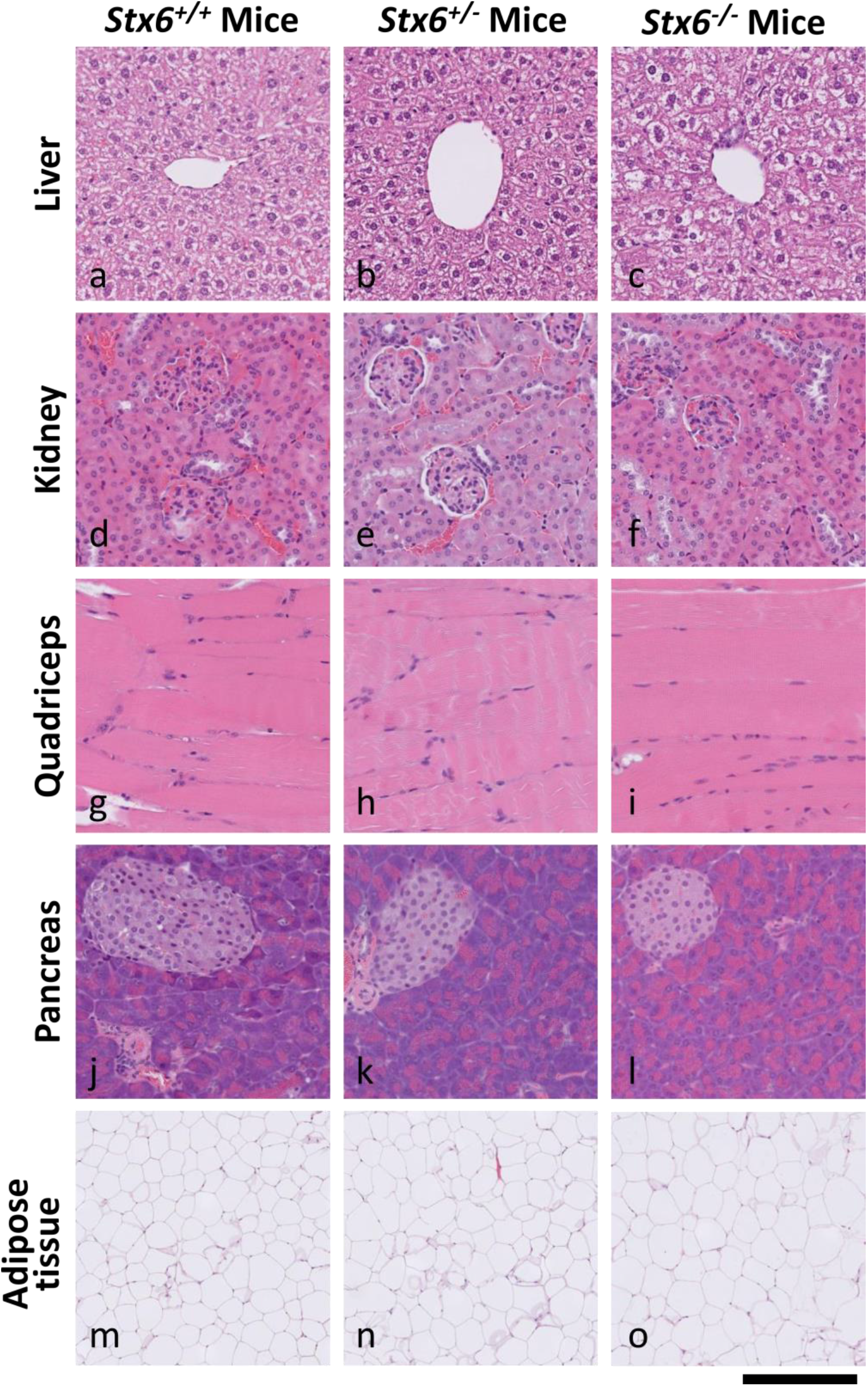
Multi-organ H&E staining shows expected tissue architecture and appearance in Stx6^-/-^ and Stx6^+/-^ mice which lack consistent tissue-intrinsic pathologies. The following organs were harvested from Stx6^+/+^ (n = 5), Stx6^+/-^ (n = 5) and Stx6^-/-^ (n = 4) female mice at 56-61 weeks of age: liver, pancreas, kidney, skeletal muscle and white adipose. These were formalin fixed, processed to paraffin and underwent H&E staining, revealing no consistent differences between genotypes. Scale bar corresponds to 120 µm (a-l), and 500 µm (m-o).

Upon generation, *Stx6*^-/-^ mice were subject to the standard IMPC phenotyping program at MRC Harwell(21) (www.mousephenotype.org). This analysis suggested alterations in some blood clinical chemistry parameters including increased blood urea nitrogen in early adult females and increased circulating alkaline phosphatase in early adult males, but these findings were not replicated in our independent sample (P > 0.01) (**Error! Reference source not found., Error! Reference source not found.**). The lack of replication or significance of these phenotypes in other sexes or life stages indicates these perturbations are not replicable or generalisable. In a pilot study, aged mice were examined on a range of behavioural and motor tasks, including nest building, rotarod, grip strength, pellets burrowed and weight of food consumed, with *Stx6*^-/-^ and *Stx6*^+/-^ (n = 5 per group) broadly comparable with wild-type animals (the study was not powered to detect subtle differences in behaviours). Overall, reduction in *Stx6* expression did not lead to a detectable deleterious phenotype in these animals.

### Mice with genetic reduction of *Stx6* have differing incubation periods when infected with two mouse-adapted prion strains

To determine the effect of *Stx6* expression on the incubation period of prion disease, *Stx6*^-/-^, *Stx6*^+/-^ and *Stx6*^+/+^ female mice were inoculated intracerebrally with two mouse-adapted scrapie prion strains, RML and ME7, or PBS as a vehicle control. An overall log-rank test of the association of disease incubation period with *Stx6* genotype demonstrated a significant relationship in both RML (P = 0.0049, **Fig 3A**) and ME7 (P < 0.0001, **Fig 3C**) inoculated animals, suggesting *Stx6* expression modulates the incubation period of prion disease with both prion strains.

**Fig 3:**
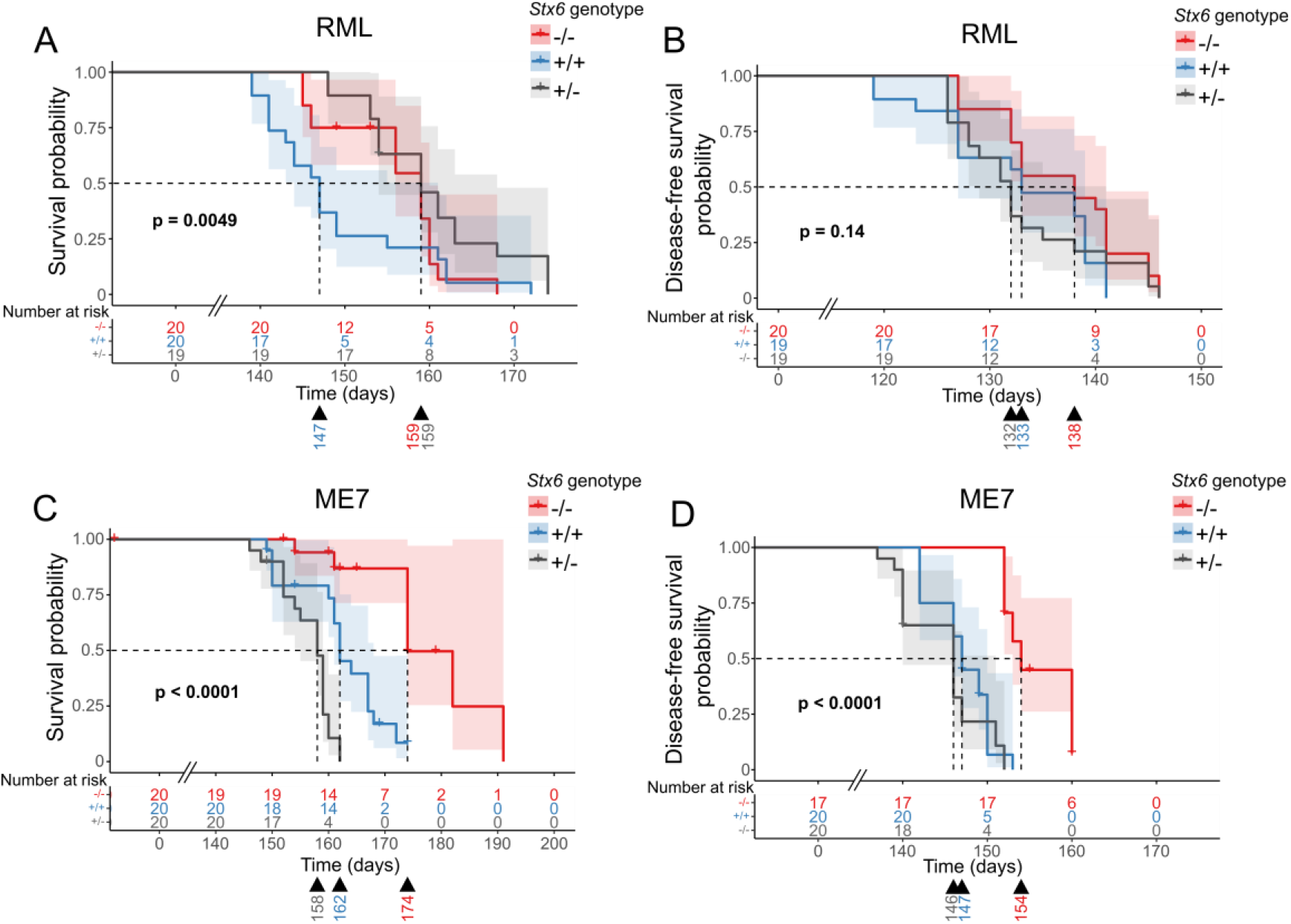
Incubation periods in mice with Stx6 reduction following intracerebral inoculation with RML and ME7 mouse-adapted scrapie prions. (**A**-**D**) Kaplan-Meier curve of survival probability following intracerebral inoculation of Stx6^-/-^ (red), Stx6^+/-^ (grey) and Stx6^+/+^ (blue) C57BL/6N mice (n = 20 per group) with 1% brain homogenate from (**A**,**B**) RML and (**C**,**D**) ME7 infected C57BL/6N mice. (**A, C**) Survival probability until animals were culled due to scrapie sickness shows a 12-day increase in median incubation period for (**A**) both Stx6^-/-^ and Stx6^+/-^ mice following RML inoculation and (**C**) for Stx6^-/-^ mice only following ME7 inoculation relative to Stx6^+/+^ mice. (**B**,**D**) Disease-free survival probability (time until onset of the first scrapie symptom) (**B**) does not show a significant association with Stx6 genotype following RML-inoculation but a nominal 5-day increase in median incubation period in Stx6^-/-^ mice compared to (**D**) a 7-day increase in Stx6^-/-^ mice inoculated with ME7, relative to Stx6^+/+^ mice. P-values indicate statistical associations with group determined using a log-rank test. Median incubation period shown below each plot. Crosses indicate censored animals.

To quantify these effects and define associations with specific genotypes, a secondary analysis was performed looking at the differences between partial and complete genetic reduction in *Stx6* expression for each strain (**Table 1**). Following inoculation with RML prions, the incubation period in *Stx6*^-/-^ and *Stx6*^+/-^ mice differed by 12 days relative to wildtype animals although statistical significance was only reached for *Stx6*^+/-^ animals (*Stx6^+/+^* = 147 days [144 – 161] (median [95% confidence interval]), *Stx6*^+/-^ = 159 days [154 – 168], *Stx6^-/-^*= 159 days [156 – 160]) (**Fig 3A**). Furthermore, as a number of animals were excluded from analysis due to early loss from intercurrent illness (especially for ME7 inoculated animals; see Materials and Methods and **Error! Reference source not found.**), the time from inoculation until onset of the first scrapie symptom was analysed (previously used as an alternative measure of incubation period(22)). There was no significant association of *Stx6* genotype with incubation period by this measure following RML inoculation (P = 0.14, **Fig 3B**), although there was still a modest, nominal increase in incubation period in *Stx6*^-/-^ animals compared to *Stx6*^+/+^ (*Stx6^+/+^* = 133 days [127 – 139] vs *Stx6^-/-^* = 138 days [133 – 141]; *Stx6*^+/-^ = 132 days [129 – 141]). Although this analysis allows for greater inclusion of animals who were succumbing to the disease, identification of the first neurological symptom is typically a more variable assessment which likely underpins the differences between these two analyses.

**Table 1:** Summary of analysis for association of Stx6 genotype with disease incubation periods following inoculation with RML and ME7 prions.

|  |  | **p\*** | **Group** | **N start** | **N events** | **Median (days)** | **Lower 95% CI** | **Upper 95% CI** | **Diff. (vs WT) (days)** | **P (vs WT)\*\*** |
| --- | --- | --- | --- | --- | --- | --- | --- | --- | --- | --- |
| **Scrapie Clinical Diagnosis** | **RML** | **P = 0.0049** | **WT** | 20 | 19 | 147 | 144 | 161 |  |  |
|  |  |  | **HET** | 19 | 18 | 159 | 154 | 168 | **+12** | 0.0047 |
|  |  |  | **KO** | 20 | 16 | 159 | 156 | 160 | **+12** | 0.17 |
|  | **ME7** | **P < 0.0001** | **WT** | 20 | 16 | 162 | 161 | 168 |  |  |
|  |  |  | **HET** | 20 | 19 | 158 | 155 | 160 | **-4** | 0.00040 |
|  |  |  | **KO** | 20 | 7 | 174 | 174 | NA | **+12** | 0.00075 |
| **First scrapie symptom** | **RML** | **P = 0.14** | **WT** | 19 | 19 | 133 | 127 | 139 |  |  |
|  |  |  | **HET** | 19 | 19 | 132 | 129 | 141 | **-1** | 0.86 |
|  |  |  | **KO** | 20 | 20 | 138 | 133 | 141 | **+5** | 0.16 |
|  | **ME7** | **P < 0.0001** | **WT** | 20 | 18 | 147 | 146 | 150 |  |  |
|  |  |  | **HET** | 20 | 19 | 146 | 140 | 151 | **-1** | 0.29 |
|  |  |  | **KO** | 17 | 14 | 154 | 153 | 160 | **+7** | 1.20 x 10<sup>-7</sup> |

Summary statistics for incubation periods measured as time until animals were culled once diagnostic clinical signs were noted (top) or onset of first scrapie-associated symptom (bottom) following intracerebral inoculation with 1% RML or ME7 infected C57BL/6 brain homogenate by Stx6 genotype (KO: -/-; WT: +/+; HET +/-). Median incubation period in days listed with corresponding 95% confidence intervals. \*P-value from overall log-rank test of Stx6 genotype with disease incubation period. \*\*P-value from a pairwise log-rank test for association of each group with relative WT controls.

In ME7-inoculated animals, incubation period also differed by 12 days with complete loss of *Stx6* compared to wildtype animals (*Stx6^+/+^* = 162 days [161 – 168]) vs *Stx6^-/-^* = 174 days [174 – NA], **Fig 3C**), however there was no increase seen with heterozygous *Stx6* expression with this strain (*Stx6*^+/-^: 158 days [155 – 160]). In the analysis of time until onset of first neurological symptom, there was still a significant association of *Stx6* genotype (P < 0.0001, **Fig 3D**) with a 7-day increase in *Stx6*^-/-^ mice compared to *Stx6*^+/+^ animals (*Stx6^+/+^* = 147 days [146 – 150] vs *Stx6^-/-^*= 154 days [153 – 160]). Again however there was no effect of partial *Stx6* expression loss on this prion strain (*Stx6^+/-^* = 146 days [140 – 151]).

Although some of the post-hoc pairwise comparisons demonstrating benefit with genetic reduction of syntaxin-6 did not reach statistical significance (**Table 1**), the independent, repeated observations of extended incubation period in *Stx6* knockout animals with two distinct prion strains supports a general pathological role of expression of this gene in prion disease progression. To explore whether *Stx6* exerts a selection effect for propagation of distinct prion strains, we performed western blotting on end stage brain homogenates. This demonstrated no change of PrP^Sc^ types with modified *Stx6* expression in either RML- or ME7-inoculated animals as expected (**Error! Reference source not found.**).

### Neuropathological features and *Stx6* expression

Prion disease is typically characterised by histological features of spongiform vacuolation, neuronal loss and astrocyte proliferation(23), along with deposition of PrP aggregates in the brain and a strong microglia response, which can be identified upon brain biopsy in patients, or on post-mortem histopathological analysis of the brain in patients and animals(24). Immunohistochemical staining with an anti-PrP antibody detected PrP deposits at the end-stage of the disease following intracerebral inoculation with RML and ME7 prions in all animals (7-10 mice analysed per group, see Materials and Methods) (**Fig 4**). With both prion strains, disease-associated PrP can be detected with a diffuse staining pattern in grey matter areas. In ME7-inoculated animals there was additional widespread deposition of micro-plaques as previously described(24). At this resolution and time-point there was no difference evident in appearance, extent or localisation of PrP aggregates between *Stx6* genotypes.

**Fig 4:**
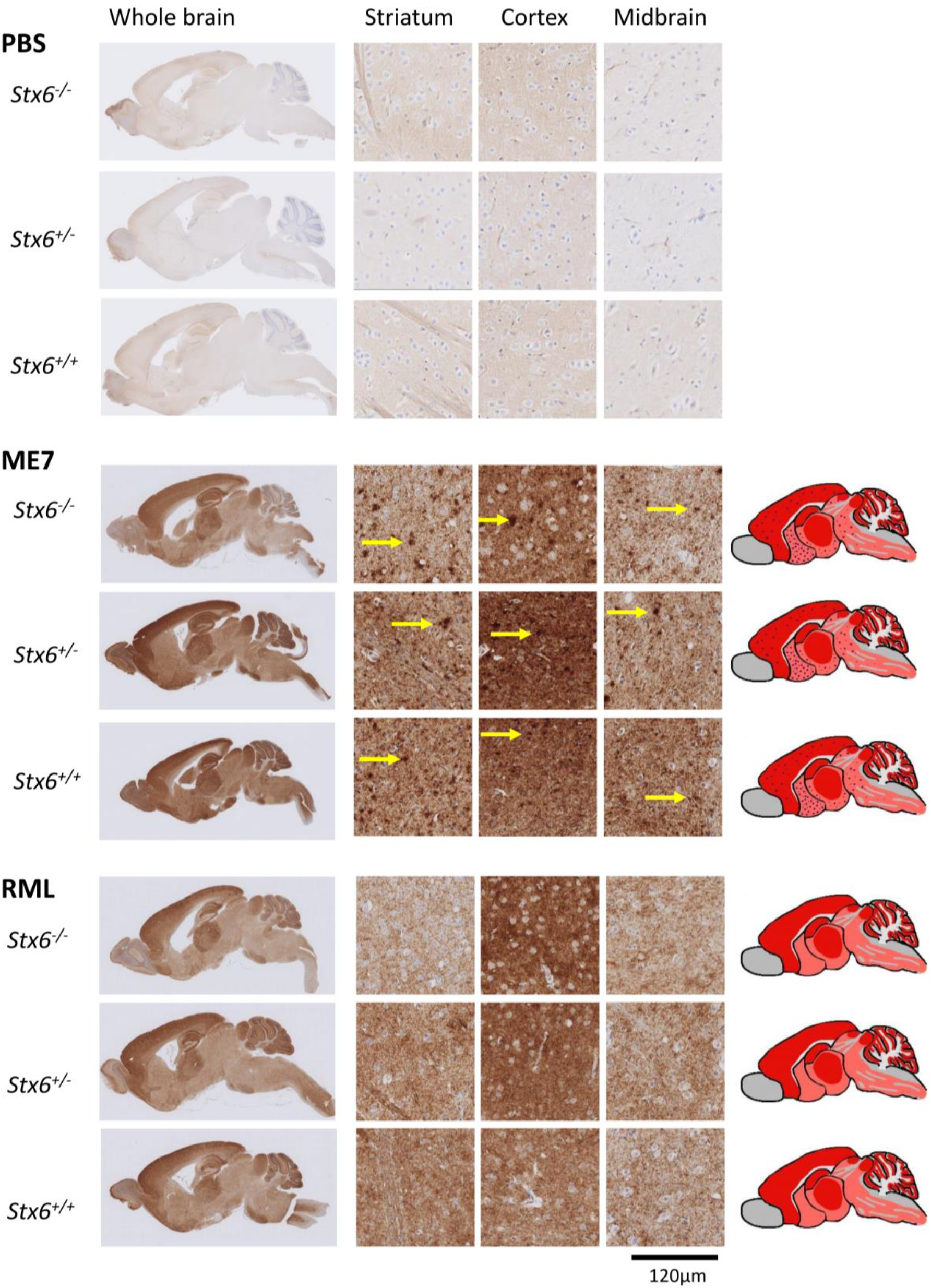
Knockout of Stx6 does not alter PrP deposition in prion infected mice. Immunohistochemistry using anti-PrP antibody ICSM35 in whole brain (left) with schematic depiction of PrP deposition (pink shading: moderate PrP deposition; red shading: intense PrP deposition; red dots: PrP micro-plaques) or representative images from striatum, cortex and midbrain (middle) from Stx6^-/-^, Stx6^+/-^ and Stx6^+/+^ mice inoculated with PBS (top) or ME7 (middle) and RML (bottom) mouse-adapted scrapie prions, shows expected PrP deposition in all prion-inoculated samples at end-point, with widespread diffuse staining for both strains and additional presence of micro-plaques with ME7 (yellow arrows). There was no evidence of spontaneous PrP deposition in control animals or gross difference evident between genotypes.

Considering this was the first characterisation of this mouse strain, we also assessed whether manipulation of this genetic risk factor alone was sufficient to cause neuropathological changes in the absence of prion infection. Knockout of *Stx6* did not cause detectable deposition of disease-associated PrP in PBS-inoculated controls.

To determine whether there were any differences in spongiform vacuolation with *Stx6* expression at endpoint, haematoxylin and eosin (H&E) staining was used to identify morphological changes in the tissue. This analysis revealed the presence of distinct small, round vacuoles within the neuropil in multiple brain regions including the striatum, cortex and midbrain in prion-inoculated animals, reflecting the expected spongiform change associated with these mouse prion strains (**Fig 5**). This analysis also showed the typical neuronal loss within the hippocampus following ME7 inoculation. No changes were identified in controls demonstrating knockout of *Stx6* alone does not induce pathological changes.

**Fig 5:**
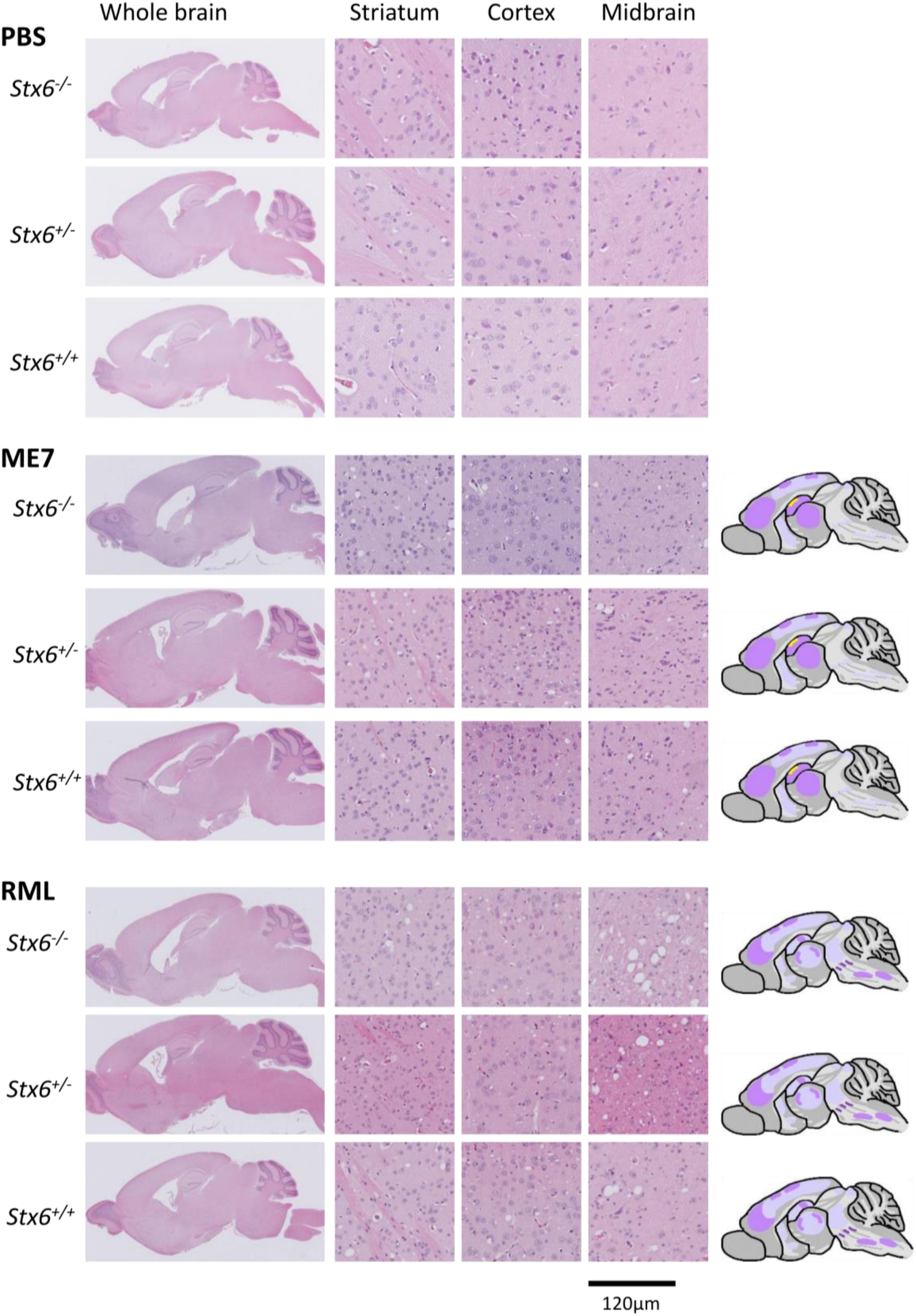
Neuronal loss and spongiform change were comparable in end-stage Stx6^-/-^, Stx6^+/-^ and Stx6^+/+^ mice. Haematoxylin and eosin (H&E) staining of whole brain (left) with schematic depiction of spongiosis (right; light purple shading: widely dispersed spongiosis; darker purple shading: intense focal spongiosis; yellow shading: neuronal loss) and representative images from striatum, cortex and midbrain (middle) from Stx6^-/-^, Stx6^+/-^ and Stx6^+/+^ mice inoculated with PBS (top) or ME7 (middle) and RML (bottom) mouse-adapted scrapie prions shows expected spongiform pathology in all samples at end-point, including hippocampal neuronal loss with ME7, with no gross differences evident between genotypes. No neuropathology was evident in PBS-inoculated controls.

Taken together, these results provide further support for maintenance of prion strain type in mice with genetic reduction of syntaxin-6, with no clear differences in PrP deposition patterns and spongiform vacuolation between genotypes.

### Possible increased inflammatory response with reduced *Stx6* expression

As widespread activation and proliferation of both astrocytes and microglia is a hallmark of both human and animal prion disease, which progresses with disease duration, immunohistochemical staining was performed to detect activated astrocytes and microglial cells allowing quantification of any modulatory effects *Stx6* expression may have on neuroinflammatory phenotypes in prion disease. Furthermore, control animals were analysed to determine if *Stx6* knockout alone causes any inflammatory phenotypes.

Immunohistochemical staining of reactive astrocytes with an anti-GFAP antibody confirmed the expected widespread astrocyte proliferation, with particular intensity in the thalamus and hippocampus as previously described with these prion strains(25) (**Fig 6A**). Quantification of mean percentage area shows >70% immunoreactivity in brain tissue under all prion-inoculated conditions (ME7: *Stx6^-/-^* = 82.3% ± 0.820 (mean ± SEM), *Stx6^+/-^*= 71.7% ± 0.870, *Stx6^+/+^* = 70.7% ± 2.15; RML: *Stx6^-/-^* = 79.2% ± 1.06, *Stx6^+/-^* = 82.6% ± 1.20, *Stx6^+/+^* = 78.6% ± 1.06) compared to ∼40% in PBS-inoculated animals (PBS: *Stx6^-/^* = 42.4% ± 1.04, *Stx6^+/-^* = 44.3% ± 0.630, *Stx6^+/+^* = 43.7% ± 0.478) (**Fig 6C**). This highlighted a ∼10% increase in astrocyte area in *Stx6*^-/-^ mice inoculated with ME7 relative to both *Stx6*^+/-^ (P_adj_ = 1.70 × 10^-3^) and *Stx6*^+/+^ animals (P_adj_ = 7.57 × 10^-4^) after adjusting for age of death to take into account the later tissue collection for the *Stx6*^-/-^ mice with extended incubation periods. Of note, there was no significant association between age and GFAP area stained. This highlights a potential protective mechanism for reduced *Stx6* expression in prion diseases, for example if astrocytes are acting to maintain neuronal function.

**Fig 6:**
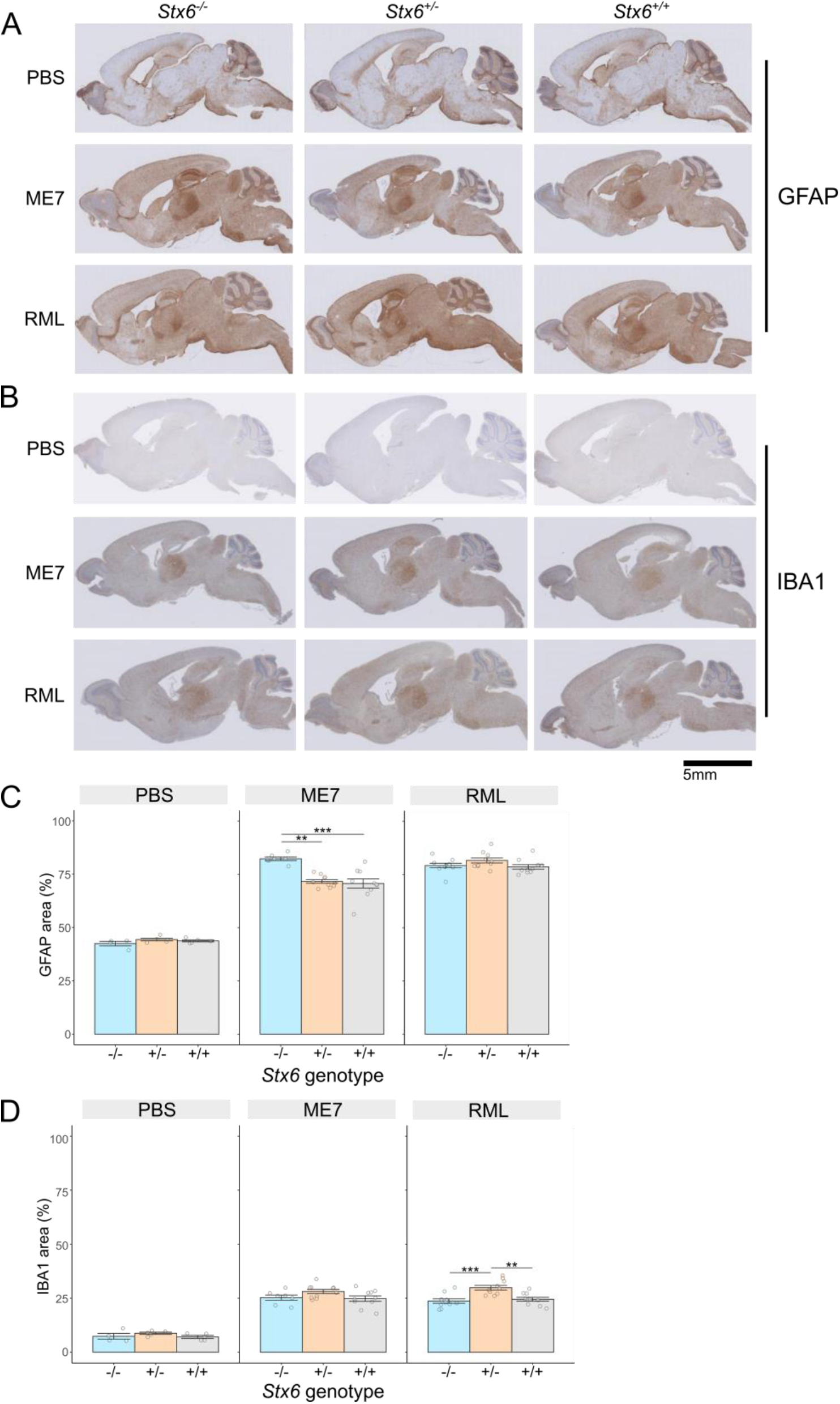
Variable effect of Stx6 genotype on neuroinflammation in prion disease. (**A, B**) Example images of immunohistochemistry in Stx6^-/-^, Stx6^+/-^ and Stx6^+/+^ mice inoculated PBS control or ME7 and RML mouse-adapted scrapie prions with (**A**) anti-GFAP antibody to measure reactive astrocytes and (**B**) anti-Iba1 antibody to measure microgliosis, shows expected neuroinflammation in prion-infected animals, including expected intense staining in the thalamus with both antibodies and intense GFAP staining in the hippocampus. (**C, D**) Quantification of percentage area stained with (**C**) anti-GFAP and (**D**) anti-Iba1 antibody demonstrates a ∼10% increase in astrocyte area in ME7-inoculated Stx6^-/-^ animals and a ∼5% increase in microglia area in Stx6^+/-^ mice in RML-inoculated animals (linear regression model adjusted for age of death; ** P < 0.01, *** P < 0.001).

Immunostaining with an anti-Iba1 antibody to detect microgliosis demonstrated the expected visible microglia activation following inoculation with both RML and ME7 prions, occupying 25-30% of the whole brain area in all disease conditions (ME7: *Stx6^-/-^*= 25.3% ± 1.20, *Stx6^+/-^* = 28.1% ± 0.972, *Stx6^+/+^* = 24.8% ± 1.26; RML: *Stx6^-/-^* = 23.7% ± 1.03, *Stx6^+/-^* = 29.9% ± 1.06, *Stx6^+/+^* = 24.5% ± 0.899) compared to ∼8% in control animals (PBS: *Stx6^-/-^* = 7.37% ± 1.33, *Stx6^+/-^* = 8.77% ± 0.525, *Stx6^+/+^* = 7.09% ± 0.700) (**Fig 6B, D**). Interestingly in all conditions *Stx6*^+/-^ mice demonstrated a modestly higher proportion of microglia relative to both *Stx6*^+/+^ and *Stx6*^-/-^ mice, which was significant in RML inoculated animals after adjusting for different ages at death (P_adj_ = 4.07 × 10^-3^ and P_adj_ = 1.74 × 10^-4^ respectively).

## Discussion

### Incubation period differences in mice with genetic reduction of syntaxin-6 following intracerebral inoculation with RML and ME7 mouse-adapted scrapie prions

Mouse models are a suitable way to assess the role of human risk genes as they are naturally susceptible to prion diseases, and prion-infected animals faithfully recapitulate many of the key aspects of human disease. In this study, C57BL/6N female mice with genetic reduction of syntaxin-6 showed prolonged mouse prion disease incubation periods for both prion strains tested. However, the observed extension of 12 days in prion-infected *Stx6^-/-^*animals was modest, which may reflect the moderate effect sizes for typical risk alleles detected by GWAS (*STX6* odds ratio (OR) = 1.16)(3) or potential compensatory mechanisms associated with constitutive gene knockout. Alternatively, it remains possible that modest changes in incubation periods might have resulted from imbalance in unmeasured confounding factors with genotypes being inoculated on different days, or simply chance effects. Indeed, when groups of genetically identical FVB/N were inoculated in different groups at different times, median incubation between individual groups ranged between 103 and 137 days despite intragroup variation being very low(26), substantially greater variation than what was observed between different arms of this study. The modest differences in incubation time observed therefore need cautious interpretation.

Although PrP^C^ levels were comparable between the genotypes suggesting a direct interaction of *Stx6* on *Prnp* expression was not the driving force behind the observation, we cannot exclude subtle effects on the expression or distribution of PrP^C^ and it remains possible, even likely, that *Stx6* modifying effects are exerted through altered trafficking of PrP^C^ or disease-associated forms of the protein. Furthermore, considering the syntaxin-6 risk effect was identified for sCJD, such an effect may not be sufficient to substantially modulate incubation times in a prion-infection paradigm using 1% infected brain homogenate with the high infectious prion dose administered being substantially different from a disease-initiating event in sporadic disease. Therefore, future experiments using lower titres are warranted to explore this further.

Although we recently identified risk variants associated with sCJD at the *STX6* locus, proximity of risk SNPs to genes is suggestive but not sufficient to establish causality. Integration of GWAS variants with eQTL data provided evidence *STX6* expression is associated with human disease risk. This *in vivo* data is therefore in keeping with the hypothesis derived from GWAS discovery that increased *STX6* expression promotes prion disease pathogenesis. Importantly however, prion transmission studies model a form of acquired prion disease, raising the possibility that syntaxin-6 plays a role across different aetiologies of prion disease and across species.

### Assessment of prion-related phenotypes

No differences in neuronal loss and the extent or distribution of PrP pathology were noted between *Stx6^+/+^, Stx6^+/-^* and *Stx6^-/-^* mice. This indicates that *Stx6* expression does not alter disease progression through a gross effect on disease pathology. However, it should be noted that this was assessed in endpoint animals where neuropathology was severe and quantification of these was not possible due to processing artefacts, which may have masked more subtle differences. Furthermore, ages of mice were not matched, as *Stx6*^-/-^ mice generally survived longer before culling than the wild-type controls.

In contrast, quantitative comparisons of gliosis between the genotypes revealed some subtle differences. Interestingly a ∼10% increase in astrocyte count was seen in ME7-infected *Stx6^-/-^* mice and a ∼5% increase in microglia count was seen in RML-infected *Stx6^+/-^* mice relative to *Stx6*^+/+^ animals. Furthermore, in all conditions *Stx6*^+/-^ mice demonstrated a modestly higher proportion of microglia relative to both *Stx6*^+/+^ and *Stx6*^-/-^ mice, which may reflect a dose-dependent effect of *Stx6* expression on microglia function. To exclude the possibility that these differences were driven by different ages of animals at endpoint, we included age at death in days as a covariate in the regression analysis with no significant effect found. Future dissection of time points across the prion incubation period may reveal additional differences in neuropathology or other readouts of disease progression.

Considering syntaxin-6 itself has been proposed as a particularly promiscuous SNARE protein with a multitude of cell-type specific functions, it is certainly plausible that the pathological role of syntaxin-6 is mediated through non-neuronal cell types. Taken together, these results suggest a possible role for syntaxin-6 in the supporting cells of the brain, fitting with the increasing acceptance that prion disease pathobiology is driven by the interaction of multiple cell types and non-cell autonomous mechanisms(27, 28).

### Targeting syntaxin-6 as a therapeutic strategy

Human genetic evidence increasingly supports successful drug development programmes(29). The identification and proposal of *STX6* as a risk gene through GWAS therefore provokes research about therapeutic potential, which is in part supported by this *in vivo* study. Importantly for therapeutic utility, *Stx6^-/-^* mice are viable and fertile with no gross neurological or physiological impairments. Although the IMPC have recorded some potential metabolic and behavioural alterations in *Stx6^-/-^*mice, these were not reproducible across life stages and gender, and were not replicated in our mouse cohort. However, the very modest extension in incubation period with full knockout of syntaxin-6 does raise the question of whether partial reduction in syntaxin-6 levels in established disease would result in a meaningful clinical benefit. Further work involves determining the efficacy of targeting *Stx6* during the disease course. Furthermore, the pleiotropic role of syntaxin-6 across multiple neurodegenerative diseases, including PSP and AD, raises the potential that *STX6*-targeting therapies could have wider applicability to other neurodegenerative diseases, warranting further investigation.

### Potential mechanistic roles of syntaxin-6 in prion disease pathology

As a key intracellular trafficking protein, it is plausible that modified syntaxin-6 function alters the transport of PrP^C^ and/or disease-associated prions throughout the cell and several possible mechanisms are plausible. Considering syntaxin-6 is thought to be involved in the recycling of proteins from endosomes(30, 31), knockout of *Stx6* could conceivably result in reduced transport of disease-associated PrP away from these structures and thus increased degradation in the lysosomal pathway, with obvious benefits for prion load. Alternatively a similar hypothesis could be proposed for transport of toxic species in prion diseases. Recent work indicated a role of syntaxin-6 in the kinetic formation of toxic PrP structures *in vitro*, however whether a direct protein interaction occurs *in vivo* is unclear(32). Finally, it is plausible the effect of modified syntaxin-6 function has more widespread implications for cellular function, which may indirectly affect prion disease biology.

## Materials and Methods

### Ethical approval

Work was performed under approval and license granted by the UK Home Office (Animals (Scientific Procedures) Act 1986), which conformed to UCL institutional and Animal Research: Reporting of In Vivo Experiments (ARRIVE) guidelines. Experiments were approved by the MRC Prion Unit Animal Research Scientific Committee.

### Mice

*Stx6*^+/-^ mice (C57BL/6NTac-*Stx6*^em1(IMPC)H^/H) were generated at MRC Harwell and crossbred to establish homozygous *Stx6*^-/-^ and *Stx6*^+/+^ lines, which were used to populate the experimental cohorts. *Stx6^+/-^* litters were obtained through crossbreeding of these two lines.

### Genotyping

DNA was extracted from standard ear biopsies. The presence of a 108 bp deletion in *Stx6* was determined using two PCR reactions with the following primer combinations with GoTaq G2 Hot Start Polymerase (Promega): PCR 1 forward 5’-CGATCTGTGAGACTCATCGGG and reverse 5’-GGGAGTCCTAACACCACCTTC, PCR 2 forward 5’-CCTGACTCTCTGATAGCCAC and reverse 5’-ACAAAACCAAAGCCTGCAC. PCR reactions were analysed by agarose gel electrophoresis and image capture was performed on a Universal Hood II Gel Doc System (Bio-Rad).

### Western blot of mouse brain homogenate for syntaxin-6

Brains from mice at 61-65 days were dissected on the sagittal plane and flash frozen. Following weighing, samples were ribolysed in Dulbecco’s phosphate-buffered saline (DPBS) with ceramic homogenisation beads (Bertin Technologies) at max speed for 105 sec to produce 10 and 20% homogenates were stored at -80° C until use.

Homogenates were diluted in DPBS with 4X SDS sample buffer (250 mM Tris base, 40% glycerol, 8% SDS, 0.04% bromophenol blue) and boiled at 95°C for 5 min. 35 µg of total protein was loaded onto a 4-12% (w/v) Bis-Tris polyacrylamide gel (Invitrogen) and electrophoresed before being electroblotted to a nitrocellulose membrane. Membranes were blocked in Odyssey Blocking Buffer for 1 h at room temperature (RT), before incubation with anti-syntaxin-6 (1:500 clone C34B2 (Cell Signalling; 2869S) or clone 3D10 (Abcam; Ab12370)) overnight at 4°C. Membranes were washed with 0.05% Tween-20 in phosphate buffered saline (PBST) for 3 × 5 min with agitation and incubated with anti-β actin (1:1000 rabbit polyclonal (Sigma Aldrich; A2066) or 1:5000 mouse monoclonal (Sigma Aldrich; A5441)). After washing, membranes were probed with fluorophore-conjugated secondary antibodies (IRDye 800CW Donkey anti-rabbit IgG and IRDye 680RD goat anti-mouse IgG both at1:4000) for 1 h at RT. Membranes were washed again and imaged with Odyssey Gel Documentation System (LI-COR; Model 9120).

### ELISA for PrP^C^ expression

Brain homogenates were initially diluted to 3.5 mg/ml total protein in PBS to a final volume of 40 μl. 5 μM AEBSF and 2.5% w/v SDS was added to each sample and boiled at 100°C for 10 min. Samples were aliquoted and stored at -80°C until use.

A high-binding 96-well flat bottom plate was pre-coated with 100 μl of 2.5 μg/ml anti-PrP ICSM18 antibody in 0.05 M carbonate coating buffer (0.035 M NaHCO_3_, 0.015 M Na_2_CO_3_, pH 9.6) and incubated at 4°C overnight before washing x3 with 1% PBST.

240 μl pre-heated capture buffer (37°C, 30X, pH 8.4; 2% N-Lauroylsarcosyl, 2% BSA, 2% Triton X-100, 50 mM Tris base pH 8.0) was added to 8 μl sample and 50 μl added to wells in triplicate. Plates were incubated at 37°C for 1 h before washing x3 with 1% PBST. For detection of bound PrP, 100 μl of 1 μg/ml biotinylated anti-PrP ICSM35 antibody diluted in 1% PBST was added to each well, incubated at 37°C for 15 min and washed as previous. 100 μl streptavidin-HRP diluted 1:10,000 in PBST was then added and incubated at 37°C for 15 min, prior to development with QuantaBlu Fluorogenic Peroxidase Substrate Kitas per manufacturer’s instructions (Thermo Scientific). Plates were imaged using Tecan Infinite M200 plate reader with 313 nm and 398 nm excitation and emission wavelengths respectively.

### Inoculation study

Anaesthetised female *Stx6^-/-^, Stx6^+/-^* and *Stx6^+/+^* mice were inoculated intracerebrally in the right parietal lobe with 30 μl 1% (w/v) C57BL/6 mouse-adapted RML or ME7 prion-infected brain homogenate at 6-8 weeks of age (n = 20/group), as previously described(33), or with PBS (n = 5/group). Experimental arms were inoculated on different days. Each individual mouse was considered an experimental unit. Group sizes of 20 mice per experimental condition were used to provide sufficient power to detect a 5% difference in incubation periods, calculated assuming an approximate minimum incubation period of 150 days with a standard deviation of 7.5 days estimated from previously published studies with C57BL/6 mice(17, 34), whilst allowing for loss due to incurrent illness (as expected from previous experiments of similar length (unpublished data)). Animals were allocated to groups based on genotypes. The animal technicians tasked with diagnosing scrapie sickness were blind to study design.

Mice were monitored daily and culled following diagnosis with scrapie sickness according to established criteria or identification of distinct health concerns(33). Incubation period was defined as the number of days from inoculation until definite scrapie diagnosis(33), with animals censored when culled due to other reasons (health concerns/adverse events outlined *a priori* in the Project licence) or found dead (**Error! Reference source not found.**).

Disease-free incubation period was defined as the number of days from inoculation to onset of the first scrapie-associated neurological symptom, with animals censored in the previous analysis due to wet genital region included in this analysis as a previously reported symptom of scrapie sickness (data unpublished). Mice culled due to other health concerns not known to be related to prion disease remained censored from this analysis (**Error! Reference source not found.**). Animals culled prior to onset of any neurological symptoms were excluded due to missingness.

Kaplan-Meier survival curves and analyses were generated using RStudio packages “survival” and “survminer” and survival differences analysed using a log-rank test.

### Western blot for prion strain typing

Following culling, brains were removed, dissected on the sagittal plane with one hemisphere flash frozen and stored at -80°C until brain homogenate preparation. 20% (w/v) homogenates were prepared in DPBS by ribolysing with ceramic homogenisation beads (Fisherbrand Zirconium Ceramic Oxide Bulk Beads) at 6,500 rpm for 45 sec using the Hybaid Riboylser. 20% homogenates were stored at -80°C until use. 20 μL aliquots of 10% (w/v) brain homogenates were prepared in DPBS. 1 μL benzonase was added and left to incubate for 10 min at RT followed by 1 h incubation with proteinase K (PK; 50 μg/mL or 100 μg/mL final concentration for ME7 and RML respectively) at 37°C with agitation (800 rpm).

Samples were mixed with 2X Lithium Dodecyl Sulphate Sample Buffer (2x dilution NuPAGE™ LDS Sample Buffer (4X) and 5x dilution NuPAGE™ Sample Reducing Agent (10X)) with 4-(2-Aminoethyl) benzenesulfonyl fluoride hydrochloride (AEBSF) (4 mM final concentration). Samples were immediately transferred to a 100 °C heating block for 10 min. Following a 1 min spin at 21,000 × g, electrophoresis was performed on 12% (w/v) bis-tris gels (Invitrogen) with SeeBlue Prestained Molecular Weight Marker prior to electroblotting to Immobilon PVDF membrane (Millipore).

Membranes were blocked in PBST with 5% (w/v) non-fat dried skimmed milk powder and then probed with ICSM35 anti-PrP antibody (D-Gen Ltd; 1:5000 for RML) or with 6D11 anti-PrP antibody (BioLegend (808004); 1:5000 for ME7) in PBST overnight. After washing (3 × 5 min followed by 3 × 15 min) the membranes were probed with a 1:10,000 dilution of alkaline-phosphatase-conjugated goat anti-mouse IgG secondary antibody (Sigma-Aldrich (A2179)) in PBST. After washing (1 h with PBST as before and 2 × 5 min with 20 mM Tris pH 9.8 containing 1 mM MgCl_2_) blots were incubated for 5 min in chemiluminescent substrate (CDP-Star; Tropix Inc) and visualized on Biomax MR film (Kodak).

### Immunohistochemistry of prion-related neuropathology

Immunohistochemistry was performed as previously described with modifications(35). Half mouse brains were fixed in 10% buffered formal-saline, processed, paraffin wax embedded and serial sections of 5 μm nominal thickness were taken. Deparaffinised sections were investigated for abnormal PrP, microgliosis and astrocytosis on the Ventana Discovery XT automated IHC staining machine (Roche Tissue Diagnostics) as described in(36). In the PBS inoculated controls, 4 × *Stx6*^-/-^, 5 × *Stx6*^+/-^ and 5 × *Stx6*^+/+^ animals were analysed (due to one *Stx6*^-/-^ mouse lost due to incurrent illness). In the RML inoculated groups, 10 mice were analysed for each genotype. In the ME7 inoculated groups, 10 mice were analysed for *Stx6*^+/+^ and *Stx6*^+/-^ groups and 7 mice in the *Stx6*^-/-^ group (only mice culled due to scrapie sickness analysed for histology). All 20 mice were analysed for neuronal loss in each group.

Anti-PrP monoclonal antibody ICSM35 (D-Gen Ltd) was used with biotinylated polyclonal rabbit anti-mouse immunoglobulin secondary antibodies (Dako; Agilent) and Ventana proprietary detection reagents utilizing 3,3′-diaminobenzidine tetrahydrochloride as the chromogen (DAB Map Detection Kit; Roche Tissue Diagnostics). Sections were treated with Discovery CC1 cell conditioning solution at 95°C for 60 min followed by a low concentration of protease (Protease 3) for 12 min prior to staining.

Anti-iba1 microglial antibody (AlphaLaboratories) or anti-GFAP antibody (Agilent) were used with biotinylated polyclonal goat anti-rabbit immunoglobulin secondary antibodies (Dako; Agilent) and Ventana proprietary detection reagents (DAB Map Detection Kit). Sections were treated with Discovery CC1 cell conditioning solution at 95°C for 60 min or a medium concentration of protease (Protease 1) for 4 min respectively prior to staining.

Conventional methods on a Gemini AS Automated Slide Stainer were used for haematoxylin staining. Positive staining controls for the staining technique were used throughout. Slides were digitally scanned on Hamamatsu NanoZoomer 360, images captured from the NDP.serve3 (NanoZoomer Digital Pathology) or NZConnect software and composed with Adobe Photoshop.

### Immunohistochemistry quantification and statistical analysis

GFAP and Iba1 immunostaining in whole brain sections was quantified using QuPath (v0.3.0) software (4-5 PBS-inoculated mice; 7-10 prion-inoculated mice). Colour deconvolution was performed to distinguish DAB staining from haematoxylin background and pixel classification used to select tissue for analysis (whole brain). Pixel classification selected regions positive for DAB staining (classifier trained on 10% total images). Area positive for DAB staining was calculated for each image and % area stained calculated relative to total tissue area analysed. Statistical differences were determined using a linear regression model with age at death included as a confounding variable in RStudio (v1.1.463) software.

### Immunohistochemistry of peripheral organs

The following organs were harvested from PBS-inoculated wildtype *Stx6^+/+^*, *Stx6^+/-^* and *Stx6^-/-^* female mice at 56-61 weeks of age: liver, pancreas, kidney, skeletal muscle and adipose tissue. Organs were fixed in 10% buffered formal-saline and processed for haematoxylin staining as previously described. Tissues were visually assessed for expected architecture integrity and for vacuolation in a semi-quantitative manner.

### Computer Tomography (CT) scanning

*Stx6^+/+^* (n = 2) and *Stx6^-/-^* (n = 2) male mice were culled at 12 weeks of age and frozen at -20 °C until CT scanning. Bone abnormalities were assessed using the Quantum x2 microCT scanner. Analysis was done on Analyse 14.0 software.

### Clinical chemistry analysis

Terminal blood was collected from 100 day old *Stx6*^+/+^ and *Stx6*^-/-^ animals by post-mortem cardiac puncture. Serum was prepared by allowing clotting for 10 mins followed by centrifugation at 2600 g at 4°C for 12 minutes. Serum was stored in polypropylene tubes at - 80°C prior to assessment of blood metabolites using the AU680 Analyser.

## Acknowledgments

We thank Florin Pintilii, Helena Costa and Fabio Argentina for further technical assistance with immunohistochemistry, Dr Jonathan Wadsworth for training support and preparation of the inoculum, the staff of the MRC Prion Unit at UCL Biological Services Facility for animal husbandry and care, Nick Kaye and Craig Fitzhugh for experimental assistance, Dr Tammy Kalber for assistance with CT scanning and Shyma Hamdan for assistance with animal genotyping. This work was funded by the UK Medical Research Council. Simon Mead and John Collinge are National Institute for Health Research Senior Investigators.

## Data availability

All data is available in **Supplementary Tables**.

