## Appendix for "Characterisation and prion transmission study in mice with genetic reduction of sporadic Creutzfeldt-Jakob Disease risk gene *Stx6*"

### Supplementary Tables

***Supplementary Table 1: Significant phenotypes identified in Stx6$^{-/-}$ mice include multiple metabolic parameters.***

| **Significant phenotype** | **Procedure** | **Parameter** | **P-value** | **Significant life stage** | **Significant sex** | **P-Value Other Gender** | **P-Value Other Life Stage** |
|---|---|---|---|---|---|---|---|
| Increased circulating alkaline phosphatase | Clinical Chemistry | Alkaline phosphatase | $6.92 \times 10^{-7}$ | Early adult | Male | $1.24 \times 10^{-4}$ | 0.19 |
| Increased blood urea nitrogen | Clinical Chemistry | Urea (Blood Urea Nitrogen - BUN) | $9.38 \times 10^{-21}$ | Early adult | Female | 0.000164 | $2.97 \times 10^{-2}$ |
| Increased circulating cholesterol | Clinical Chemistry | Total cholesterol | $1.48 \times 10^{-6}$ | Late adult | Male | 0.190 | 0.54 |
| Abnormal vibrissae morphology | Combined SHIRPA and Dysmorphology | Vibrissae - appearance | $6.84 \times 10^{-5}$ | Early adult | Male | 1 | 0.25 |
| Abnormal gait | Combined SHIRPA and Dysmorphology | Gait | $6.38 \times 10^{-5}$ | Early adult | Both | N/A | 0.55 |
| Hyperactivity | Open Field | Periphery (P) resting time and whole arena (WA) resting time | $4.27 \times 10^{-5}$ (P)<br>$8.82 \times 10^{-6}$ (WA) | Early adult | Female | 0.00573 (P)<br>0.000163 (WA) | $1.93 \times 10^{-3}$ (P)<br>0.15 (WA) |
| Decreased anxiety related response | Open Field | Percentage centre movement time | $7.83 \times 10^{-5}$ | Late adult | Female | 0.132 | $5.41 \times 10^{-4}$ |

*Phenotyping tests were performed by MRC Harwell as part of the IMPC including morphological, physiological and behavioural measurements measured at early adult (week 9-16) and late adult (week 52-59) life stages. Differences between Stx6$^{-/-}$ mice (Stx6$^{em1(IMPC)H}$) and wildtype C57BL6/N mice from the same facility analysed using a linear mixed model (continuous data) or Fisher's exact test (categorical data). Listed are all associated phenotypes below the $P < 10^{-4}$ threshold. Data and analyses downloaded from www.mousephenotype.org.* Data accessed 9th January 2023.

***Supplementary Table 2: Number of mice included in survival analyses.***

| Analysis | Prion strain | Stx6 genotype | N start | N events |
|---|---|---|---|---|
| Scrapie sickness | RML | +/+ | 20 | 19 |
| | | +/- | 19 | 18 |
| | | -/- | 20 | 16 |
| | ME7 | +/+ | 20 | 16 |
| | | +/- | 20 | 19 |
| | | -/- | 20 | 7 |
| First scrapie symptom | RML | +/+ | 19 | 19 |
| | | +/- | 19 | 19 |
| | | -/- | 20 | 20 |
| | ME7 | +/+ | 20 | 18 |
| | | +/- | 20 | 19 |
| | | -/- | 17 | 14 |

### Supplementary Figures

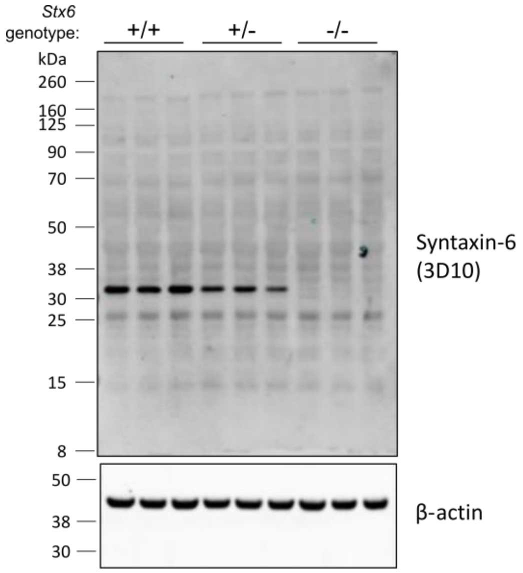


***Supplementary Fig 1: Stx6 expression in whole brain homogenate with additional anti-syntaxin-6 antibody.*** *Quantitative immunoblot of whole brain homogenates from Stx6$^{+/-}$ and Stx6$^{-/-}$ mice relative to wildtype Stx6$^{+/+}$ littermate controls demonstrates loss of primary ~32 kDa protein isoform in Stx6$^{-/-}$ mice with ~50% expression in Stx6$^{+/-}$ mice.*

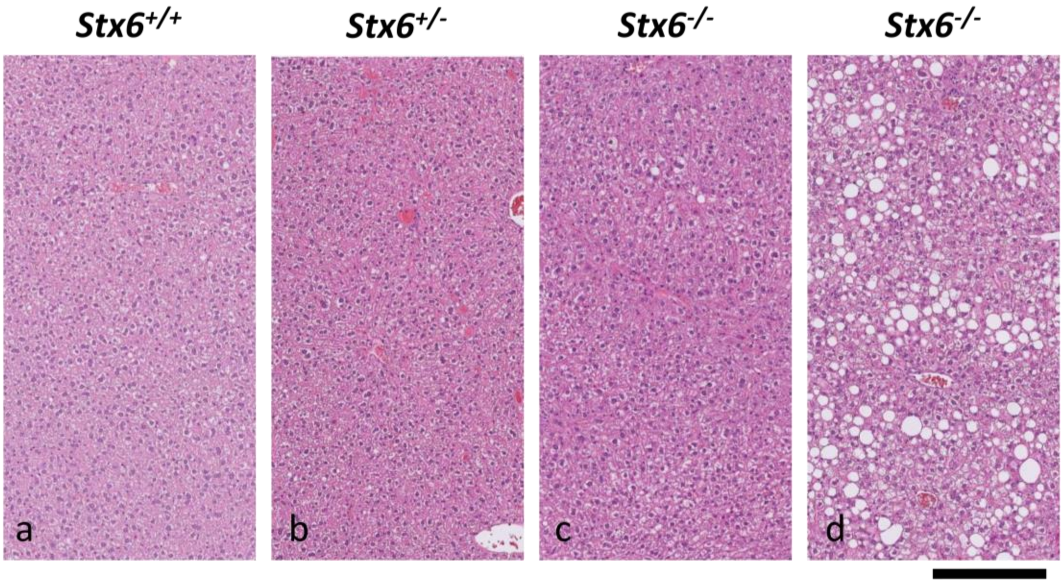


693

694 ***Supplementary Fig 2: Moderate liver steatosis in one Stx6$^{-/-}$ animal.*** *H&E staining of liver*
695 *from Stx6$^{+/+}$, Stx6$^{+/-}$ and Stx6$^{-/-}$ mice (n = 4-5 per group; a-c) shows the expected appearance*
696 *in all animals except moderate steatosis in one Stx6$^{-/-}$ animal (d). Scale bar corresponds to 120*
697 *µm.*

698

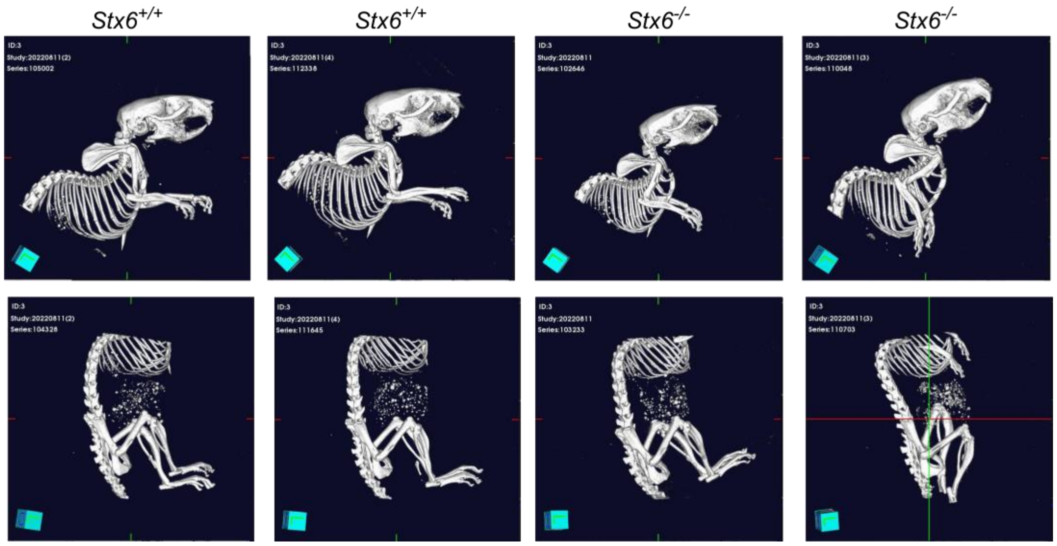


***Supplementary Fig 3. No obvious differences in skeletal structure in Stx6*$^{-/-}$ *mice compared to Stx6*$^{+/+}$ *mice.*** *Bones of Stx6*$^{+/+}$ *and Stx6*$^{-/-}$ *cadavers (n = 2/genotype; 3 months of age) were assessed by Computed Tomography (CT) scanning. Upper body (top) and lower body (bottom) were scanned separately.*

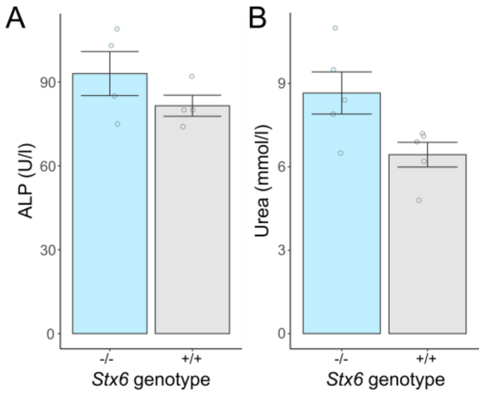


***Supplementary Fig 4: Validation of clinical chemistry phenotypes in early adult Stx6 knockout mice.*** *Analysis of significantly altered clinical chemistry parameters from IMPC analysis in serum of 100-day old (early adult) Stx6$^{-/-}$ and Stx6$^{+/+}$ mice for levels of (**A**) alkaline phosphatase (males) (ALP) and (**B**) urea (females) all P > 0.01 (Student's t-test) (mean ± SEM).*

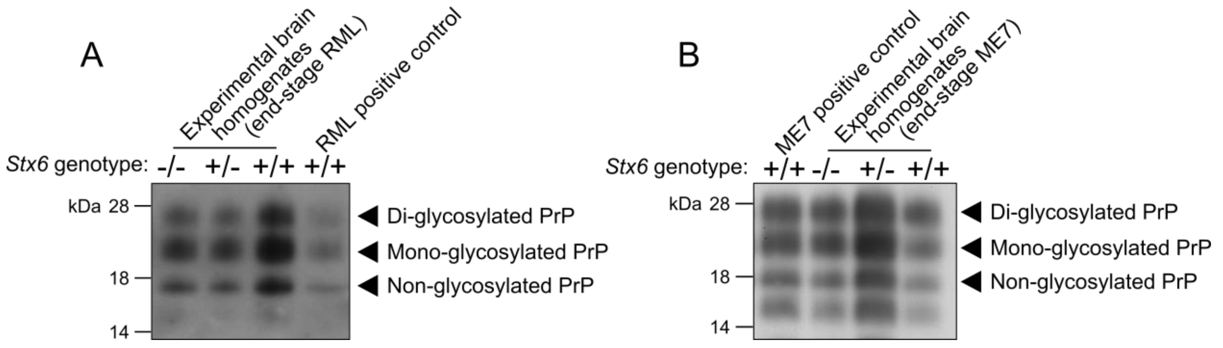


***Supplementary Fig 5: No effect of Stx6 expression on end-stage RML or ME7 prion strain type.*** *Western blot analysis of PK-resistant PrP in brain homogenates from (**A**) RML and (**B**) inoculated animals at disease end-stage shows expected electrophoretic mobility and glycosylation pattern of RML and ME7 inoculum respectively with all Stx6 genotypes.*
